# A targetable epigenetic vulnerability in PI3K/AKT inhibitor resistant cancers

**DOI:** 10.1101/2020.08.27.269613

**Authors:** Di Wu, Yuqian Yan, Ting Wei, Zhenqing Ye, Yutian Xiao, Yunqian Pan, Jacob J. Orme, Dejie Wang, Liguo Wang, Shancheng Ren, Haojie Huang

## Abstract

Acquisition of resistance to PI3K/AKT-targeted monotherapy implies the existence of common resistance mechanisms independent of cancer type. Here we demonstrate that PI3K/AKT inhibitors cause glycolytic crisis, acetyl-CoA shortage and a global decrease in histone acetylation. Also, PI3K/AKT inhibitors induce drug resistance by selectively augmenting H3K27 acetylation and binding of CBP/p300 and BRD4 proteins at a subset of growth factor and receptor (GF/R) gene loci. BRD4 occupation at these loci and drug resistant cell growth are vulnerable to both bromodomain and HDAC inhibitors. Little or none occupation of HDAC proteins at the GF/R gene loci underscores the paradox that cells respond equivalently to the two classes of inhibitors with opposite modes of action. Targeting this unique epigenetic vulnerability offers two viable strategies to overcome PI3K/AKT inhibitor resistance in different cancers.

## Introduction

Activation of receptor tyrosine kinases (RTKs) in response to extracellular stimuli, including growth factors and cytokines, enhances phosphoinositide-3 kinase (PI3K) activity, which in turn initiates a kinase cascade that subsequently phosphorylates and activates the downstream effector AKT. Active AKT plays a central role in an array of biological processes including glucose utilization, protein synthesis and cell growth and survival (Fruman *et al*, 2017; Liao & Hung, 2010; Manning & Toker, 2017). Genomic lesions activating PI3K/AKT signaling, such as the mutations in *PIK3CA, AKT* or *RAS* and loss of *PTEN, TSC* or *INPP4B*, comprise one of the most frequently mutated signaling pathways in human cancers (Engelman, 2009). Many cancer hallmarks including metabolism, survival and progression have been linked closely to the aberrant activation of the PI3K/AKT signaling pathway (Klempner *et al*, 2013).

Hyperactivation of the PI3K/AKT signaling pathway has been actively pursued as a therapeutic target in cancers. A number of PI3K/AKT inhibitors have been developed and tested under preclinical or clinical settings (Rodon *et al*, 2013). Surprisingly, PI3K or AKT inhibitor monotherapy has shown limited efficacy (Fruman & Rommel, 2014; Jansen *et al*, 2016; Nitulescu *et al*, 2016; Okkenhaug *et al*, 2016). Compared to RTK inhibitors, which usually induce dramatic therapeutic response for months or years before tumor acquires resistance (Kobayashi *et al*, 2005; Wilson *et al*, 2012), PI3K/AKT inhibitors often trigger primary resistance at clinical doses, implying that an adaptive mechanism compensates for PI3K/AKT blockage (Hopkins *et al*, 2018; Klempner *et al.*, 2013; Pan *et al*, 2017).

Feedback activation of transcription factors is implicated in the acquisition of resistance to PI3K/AKT inhibition. AKT activation results in phosphorylation and cytoplasmic localization of O class of forkhead box (FOXO) transcription factors (Huang & Tindall, 2007). Conversely, inhibition of AKT leads to FOXO protein dephosphorylation and translocation to the nucleus. Feedback expression of RTKs plays important roles in resistance to inhibitors of PI3K, AKT and HER2, and these processes are mediated, at least in part by FOXO transcription factors (Chandarlapaty *et al*, 2011; Garrett *et al*, 2011). Aberrant activation of estrogen receptor (ER) can also mediate the compensatory effects upon inhibition of PI3K in breast cancer cells (Toska *et al*, 2017), and this resistance can be attenuated by co-targeting of bromodomain and extra-terminal motif (BET) proteins including BRD4 (Stratikopoulos *et al*, 2015). Inhibition of PI3K in PTEN-negative prostate cancer cells also leads to drug resistance due to androgen receptor (AR)-dependent up-regulation of *HER2/3* kinase genes (Carver *et al*, 2011; Mulholland *et al*, 2011). Thus, feedback up-regulation of RTKs by activation of transcription factors has been implicated in the acquisition of resistance to PI3K/AKT inhibitors in cancers. However, the involved feedback-activated RTKs vary among tumors, making the precise targeting of specific RTKs in any given tumor unpredictable.

In the present study we aim to define a common mechanism and therapeutic strategy to overcome PI3K/AKT inhibitor resistance in different cancer types. Unlike previous transcription factor-centric paradigms, we identify an intrinsic epigenetic trait that is not only the major driving force for PI3K/AKT inhibitor resistance, but is also vulnerable to both bromodomain and HDAC inhibitors.

## Results

### Inevitable resistance to PI3K/AKT inhibitors independent of cancer types

Aberrant activation of the PI3K/AKT pathway due to genetic alterations such as *PTEN* mutation/deletion not only plays a causal role in tumorigenesis but is also a promising therapeutic target (Hopkins *et al.*, 2018; Jamaspishvili *et al*, 2018). Much to our surprise, we found that PTEN-negative prostate cancer cell lines PC-3 and C4-2 grew continuously after treatment with GDC-0068 (termed GDC hereafter), a highly selective AKT inhibitor which is being investigated in phase II clinical trials (de Bono *et al*, 2019; Lin *et al*, 2013; Yan *et al*, 2013) (**Fig EV1A and B**). We further challenged these cell lines with escalating concentrations of GDC over time, incrementing by 5 μM every 2 days in PC-3 cells and 1 μM every 2 days in C4-2 cells for 10 cycles. Both cell lines were trained with GDC or other PI3K (LY294002) and AKT (MK-2206) inhibitors (**Table EV1**) and their resistance to treatment was determined using MTS assays. We demonstrated that GDC-resistant (termed GDC-R hereafter) PC-3 and C4-2 cells (**Fig EV1C and D**) were also resistant to MK-2206 (**Fig 1A**). We also established MK-2206- and LY294002-resistant (termed MK-R or LY-R) PC-3 cell lines (**Fig EV1E and F**) using a similar approach and found that both were resistant to MK-2206 (**Fig 1A**).

**Figure 1.**
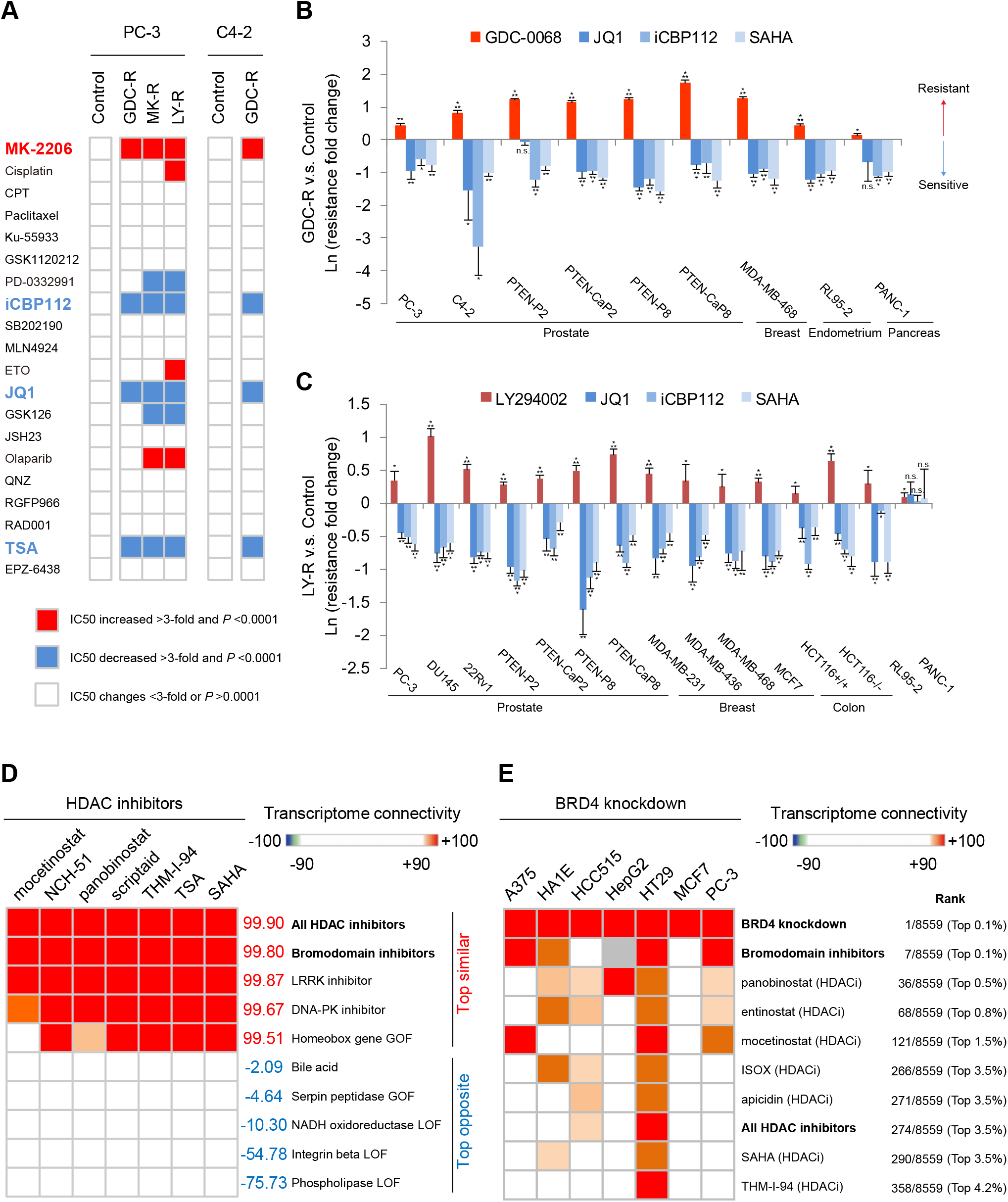
PI3K/AKT inhibitor resistant cancer cells are paradoxically hypersensitive to both bromodomain and HDAC inhibitors. A. Heatmap showing the vulnerability of AKT inhibitor GDC-0068 resistant (GDC-R) and MK-2206 resistant (MK-R) or PI3K inhibitor LY294002 resistant (LY-R) cells to compounds targeting distinct pathways (**Table EV1**). Results highlighted in red or blue in the heatmap show the significantly increased or decreased IC50 for the indicated compound in the resistant cells compared to the control cells. B, C. GDC-0068 (or LY294002)- and mock-trained cells were treated with GDC-0068 (or LY294002), JQ1, iCBP112 or SAHA (at the IC50 concentration of each compound in the control cells) for 48 hrs. The inhibitory rate in the control cells divided by that in the trained cells and the quotient was defined as the resistance fold change. The magnitude of the resistance was demonstrated as the Ln value of resistance fold change. Error bars: standard deviation (SD). Data are presented as the mean ± SD from 3 biological replicates. \**P<*0.05, \*\**P<*0.01, \*\*\**P<*0.001; n.s.: not significant. D. Transcriptome connectivity analysis shows the list of treatments that induced the most similar (or the most opposite) transcriptome effect like the pan-HDAC inhibitors in multiple cancer cell lines. The higher connectivity score indicated the closer similarity with the effect of HDAC inhibitors. The data were obtained from the Cell Connectivity Map database. E. Transcriptome connectivity analysis of 8,559 chemical or genetic interruptions, ranked by the similarity of their effects with those of BRD4 knockdown in 7 cell lines indicated. The similarities of the effects of individual HDAC inhibitors with those of BRD4 knockdown were ranked in the top 5% of all chemical or genetic interruptions examined.

Next, we surveyed GDC and LY294002 resistance in a range of cancer cell lines of different organ origins. Each of these cell lines bears one or more key mutations in the PI3K/AKT pathway genes (**Table EV2**). We identified 9 GDC-R cell lines (**Fig 1B**) and 15 LY-R cell lines (**Fig 1C**) on treatment of 16 parental cell lines with escalating doses of inhibitors. Thus, consistent with previous reports (Fruman & Rommel, 2014; Jansen *et al.*, 2016), our data showed that resistance to PI3K-AKT pathway inhibitors emerges independently of cancer type.

### PI3K/AKT inhibitor resistant cancer cells are paradoxically hypersensitive to both bromodomain and HDAC inhibitors

To explore the therapeutic vulnerabilities of these PI3K/AKT inhibitor-resistant cells, we treated resistant PC-3 and C4-2 cells with an array of chemotherapeutic drugs or tool compounds targeting various signaling pathways (**Table EV1**). Compared to control cells, resistant cells were either hyper or less sensitive to specific compounds (**Fig 1A**). Although not all positive hits were consistent across cell lines, we were able to demonstrate that GDC-, MK- and LY-resistant PC-3 and GDC-resistant C4-2 cell lines were invariably hypersensitive to BET bromodomain inhibitor JQ1, CBP/p300 bromodomain inhibitor iCBP112 and HDAC inhibitor trichostatin A (TSA) (**Fig 1A**).

Bromodomain proteins recognize acetylated histone and non-histone proteins and generally act to promote transcription (Roe *et al*, 2015), whereas HDAC proteins suppress transcription by removing histone acetyl-modifications. It seemed counterintuitive that inhibition of either bromodomain proteins or HDACs could equivalently overcome PI3K/AKT inhibitor resistance in cancer cells. To ensure these findings were reproducible, we treated GDC-resistant PC-3 cells with another CBP/p300 bromodomain inhibitor CPI-637 and another HDAC inhibitor vorinostat (SAHA), an anti-cancer chemotherapy drug. As with iCBP112 and TSA, GDC-resistant PC-3 cells were much more sensitive to both CPI-637 and SAHA in comparison to control cells (**Fig EV1G**). Most importantly, this phenomenon was not limited to PC-3 cell line since all other 8 GDC-R cell lines and 13 LY-R cell lines (except PANC-1) we identified were also hypersensitive to JQ1, iCBP112 and SAHA (**Fig 1B and C**). These data show that almost all PI3K/AKT inhibitor resistant cancer cell lines we tested are vulnerable to the inhibition of both BET and CBP/p300 bromodomain-containing proteins and HDACs.

As BET proteins, CBP/p300 and HDACs all function as important transcriptional regulators, we sought to determine the effect of bromodomain and HDAC inhibitors on gene expression in a large variety of cancer cell lines of different tissue origins. Using the Cell Connectivity Map (CMAP) (Lamb *et al*, 2006; Subramanian *et al*, 2017), an unbiased high-throughput screening method, we quantified and compared the cell transcriptome signatures under thousands of pharmaceutical or genetic interruptions. Out of more than 8,000 types of treatment, we identified compounds with HDAC inhibitor-like effects on the cell transcriptome. Surprisingly, we found that bromodomain and HDAC protein inhibitors induced very similar gene expression patterns in various cancer cell lines (**Figs 1D and EV1H**). These findings were further supported by the observation in 7 different cell lines that the similarities of the effects of HDAC inhibitors on gene expression changes with those of BRD4 knockdown were ranked in the top 5% of all treatments examined (>8,000 types) (**Fig 1E**). These unexpected results indicate that both bromodomain and HDAC inhibitors are able to overcome resistance to PI3K/AKT inhibitors regardless of cancer origin.

### PI3K/AKT inhibitors suppress glycolysis and histone acetylation

AKT activation correlates with histone acetylation level in prostate cancer (Lee *et al*, 2014). The activity of bromodomain and HDAC proteins both revolve around histone acetylation. We hypothesized that histone acetylation may play a pivotal role in the PI3K/AKT inhibitor resistance. To test this hypothesis, we assessed histone acetylation in PC-3 and C4-2 cell lines acutely treated with a panel of PI3K/AKT pathway-related inhibitors. We demonstrated that culture of cells in the absence of serum or in the presence of inhibitors of PI3K and AKT substantially decreased histone acetylation marks including total histone H3 and H3K27 acetylation (termed H3K27-ac hereafter) (**Figs 2A and EV2A**). The mTOR inhibitor RAD001 did not. In contrast, treatment with growth factors EGF, IGF and bFGF increased histone acetylation in an AKT-dependent manner (**Fig 2B**). PI3K/AKT inhibition induced down-regulation of histone H3 acetylation in all 9 GDC-resistant cell lines and in 12 of 15 LY-resistant cancer cell lines (**Fig EV2B and C**). These data suggest that there exists a common mechanism linking histone acetylation alterations and PI3K/AKT inhibitor resistance in different cancer types.

**Figure 2.**
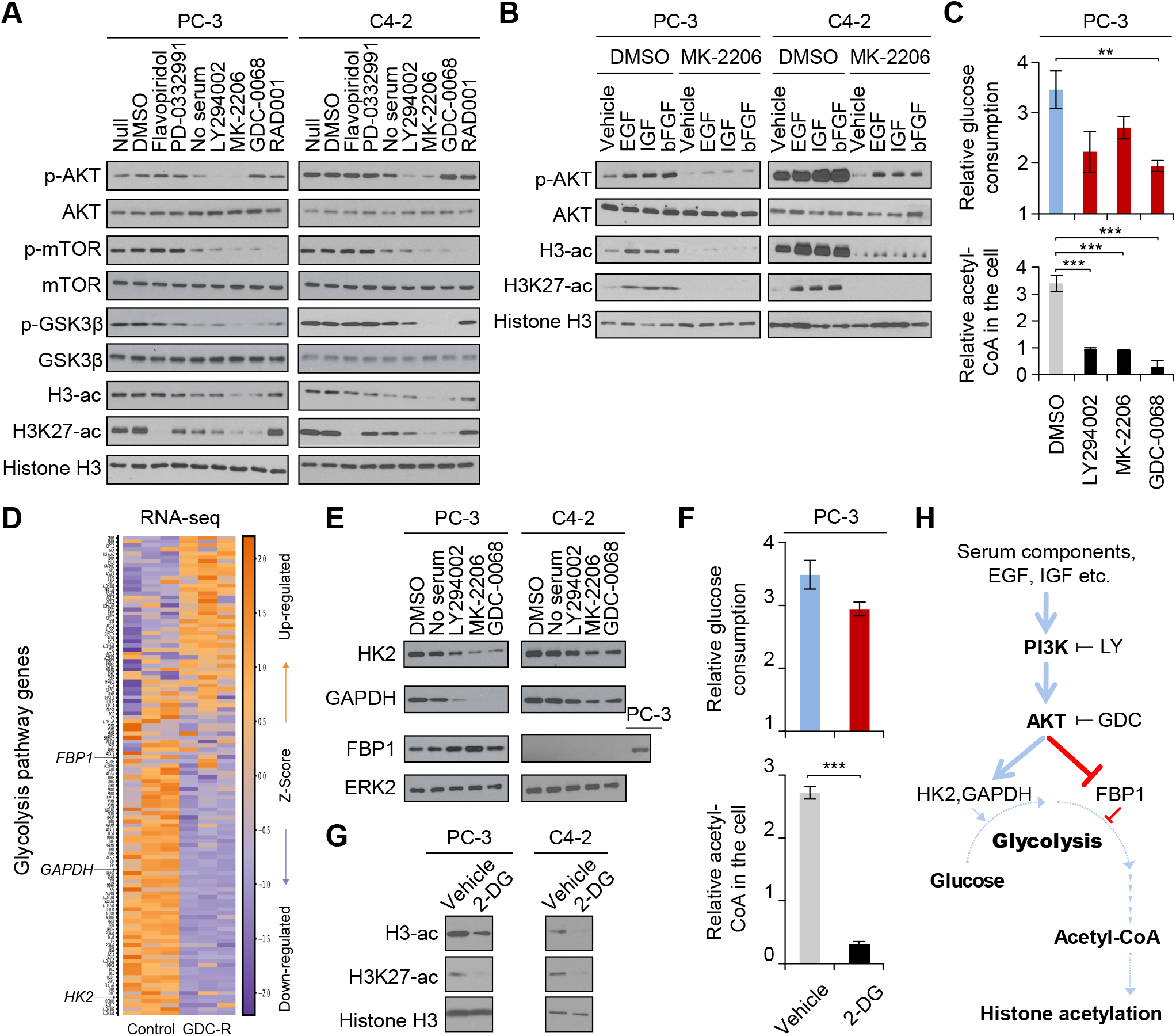
PI3K/AKT inhibitors suppress glycolysis and histone acetylation. A. PC-3 and C4-2 cells were treated with nothing (Null), vehicle (dimethyl sulfoxide, DMSO) or the indicated compounds or conditions for 24 hrs followed by western blotting. The target of each compound is listed in **Table EV1**. AKT serine 473 phosphorylation (p-AKT), mTOR serine 2448 phosphorylation (p-mTOR) and GSK3β serine 9 phosphorylation (p-GSK3β) were used to confirm the effectiveness of PI3K or AKT inhibition. B. PC-3 and C4-2 cells pre-treated with DMSO or 1 μM MK-2206 for 2 hrs were further treated with the indicated growth factors for another 24 hrs. The cell lysate and acid-extracted histone were analyzed via western blotting. C. PC-3 cells were treated with the indicated inhibitors for 48 hrs. The average medium glucose consumption rate was calculated via quantified glucose levels at the start and end points. The end point intracellular acetyl-CoA was also quantified respectively. Data are presented as the mean ± SD from 3 biological replicates. \*\**P<*0.01, \*\*\**P<*0.001. D. Heatmap shows altered expression of the genes encoding glycolytic enzymes between the GDC-resistant (GDC-R) PC-3 cells and the control cells. E. PC-3 and C4-2 cells were treated as indicated for 24hrs before harvested for western blotting. F. PC-3 cells were treated with vehicle (PBS) or 1 mM 2-Deoxy-D-glucose (2-DG) for 48 hrs. The average glucose consumption rate was calculated. The end point intracellular acetyl-CoA was also quantified. Data are presented as the mean ± SD from 3 biological replicates. \*\*\**P<*0.001. G. Cells were treated with vehicle (PBS) or 1 mM 2-DG for 48 hrs followed by western blotting. H. A hypothetical model depicting the molecular basis of PI3K/AKT inhibition-induced suppression of glycolysis, shortage of acetyl-CoA supply and decrease of histone acetylation.

On the spectrum of metabolic processes, glycolysis profoundly influences histone acetylation through acetyl-CoA levels (Moussaieff *et al*, 2015), a rate-limiting substrate of nuclear acetyltransferases (Reid *et al*, 2017). It has been suggested that AKT activation promotes glycolysis in mammalian cells (Wang *et al*, 2014). We demonstrated that PI3K/AKT inhibition not only decreased glucose consumption, but also significantly reduced acetyl-CoA level in PC-3 cells (**Fig 2C**).

To define the key factors mediating the effects of PI3K/AKT signaling on histone acetylation, we analyzed the expression of glycolytic enzyme genes in our RNA sequencing (RNA-seq) data generated from the control and GDC-resistant PC-3 cell lines (**Fig 2D**). Expression of glycolysis rate-limiting enzyme *HK2* and *GAPDH* mRNA was significantly lower in resistant cells compared to control cells (**Figs 2D and EVS2D-F**). Culture of PC-3 and C4-2 cells in the absence of serum or in the presence of PI3K/AKT inhibitors substantially decreased HK2 and GAPDH protein expression (**Fig 2E**). Similarly, pharmacological inhibition of HK2 by 2-deoxy-D-glucose (2-DG) decreased glucose consumption and intracellular acetyl-CoA level in PC-3 cells (**Fig 2F**). 2-DG treatment also decreased histone H3 acetylation in both PC-3 and C4-2 cells (**Fig 2G**).

FBP1 is another rate-limiting enzyme, but inhibits glycolysis (Cong *et al*, 2018) (**Fig EV2D**). While there was no significant difference in the *FBP1* mRNA level between the control and GDC-resistant PC-3 cell lines (**Figs 2D and EV2G**), acute treatment of PC-3 cells with PI3K/ATK inhibitors increased FBP1 protein level (**Fig 2E**). Knocking down FBP1 increased histone H3 acetylation, glucose consumption and intracellular acetyl-CoA level in PC-3 cells (**Fig EV2H-J**). Notably, expression of FBP1 protein was barely detectable in C4-2 cell line (**Fig 2E**), suggesting that the role of FBP1 in leveraging glycolysis and histone acetylation level is insignificant in C4-2 cells. Nevertheless, our data demonstrated that PI3K and AKT inhibitors induce decreased expression of HK2 and GAPDH and cell type-specific upregulation of FBP1, triggering glycolysis inhibition and global down-regulation of histone acetylation (**Fig 2H**).

### Locus-specific gain of H3K27-ac and BRD4 occupancy amid global decrease of histone acetylation in the AKT inhibitor resistant cancer cells

Given that overall histone acetylation is down-regulated in PI3K/AKT inhibitor resistant cells (**Figs 2A and EV2A**) and H3K27-ac remains low beyond 24 hours (**Fig EV3A**), we carried out chromatin immunoprecipitation sequencing (ChIP-seq) to characterize the genome-wide distributions of H3K27-ac and H3K27 trimethylation (H3K27-me3) modifications in control versus GDC-resistant PC-3 cells after 24 hours of treatment. We only observed very subtle difference in the global distribution of H3K27-me3 between control and GDC-resistant cells (data not shown). In contrast, there was a significant change of H3K27-ac distribution in GDC-resistant cells compared to control cells (**Fig EV3B**). A large number of H3K27-ac loci were lost in GDC-resistant cells, reflecting the global decrease of H3K27-ac (**Fig 3A**). Intriguingly, H3K27-ac was significantly increased at 1,332 loci in GDC-resistant cells compared to controls, and 22% and 78% of these loci were located in promoter and non-promoter regions (e.g. enhancers), respectively (**Figs 3A and EV3C**).

**Figure 3.**
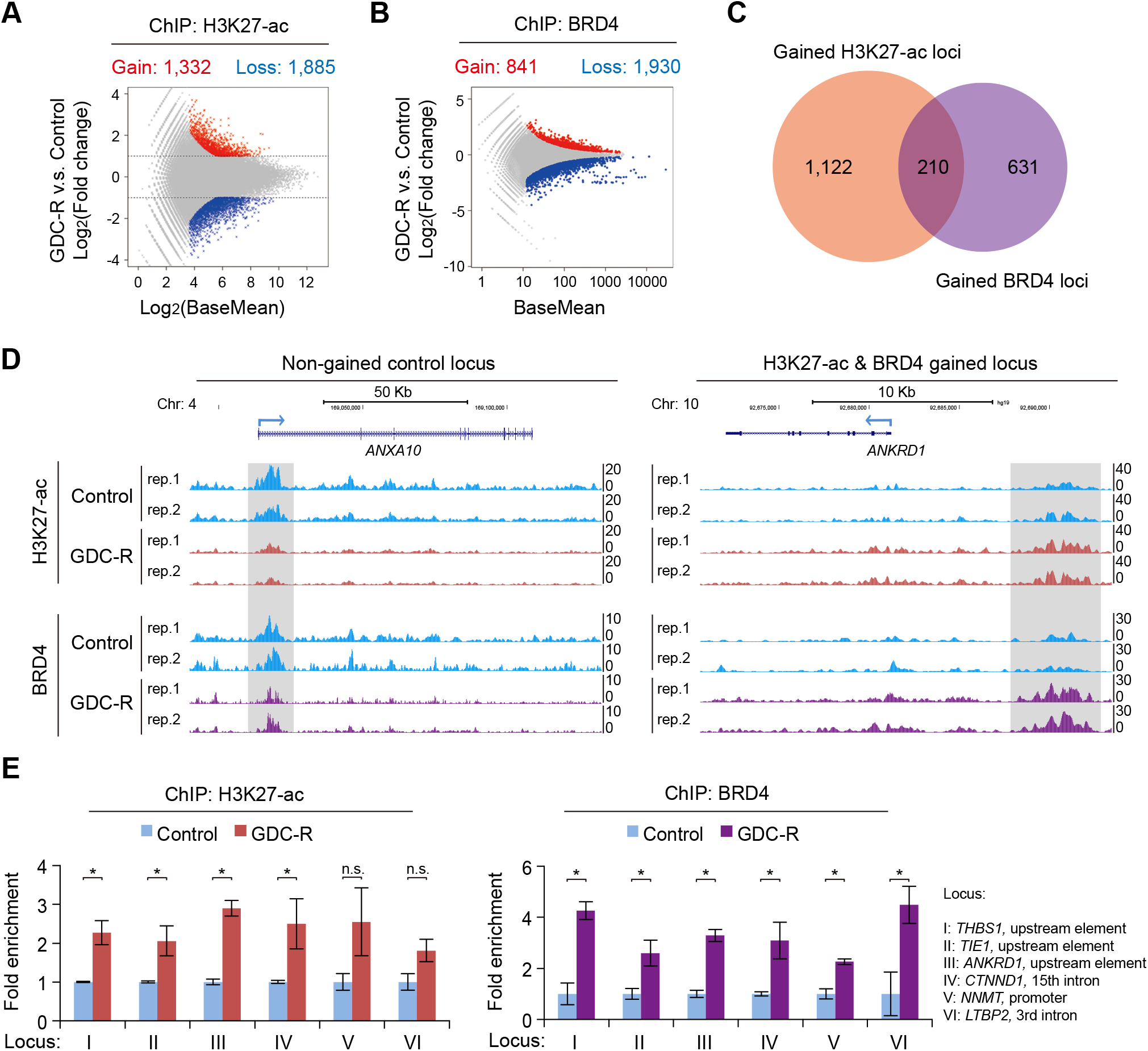
Locus-specific gain of H3K27-ac and BRD4 occupancy amid global decrease of histone acetylation in the AKT inhibitor resistant cancer cells. A, B. H3K27-ac (A) and BRD4 (B) ChIP-seq in duplicates was carried out in the control and GDC-R PC-3 cells. Each reproducible peak called from ChIP-seq reads in the resistant cells was compared with that in the control cells at the same genome location and represented as a dot in the diagram. Dashed lines indicated the thresholds that defined gained (red) or lost (blue) loci. C. Venn diagram showing the number of loci with gained H3K27-ac and BRD4 binding in the PC-3 GDC-R cells. D. Examples of the gene loci with or without gain of H3K27-ac and BRD4 ChIP-seq signal intensity (Scale, signal per 10 million reads) in the PC-3 GDC-R cells. Gray boxes, the regions with or without gain of H3K27-ac level and BRD4 binding. rep, ChIP-seq replicates. E. ChIP-qPCR analysis targeting the indicated genome loci with increased enrichment of H3K27-ac and BRD4 binding in the PC-3 GDC-R cells compared to the control cells. Primers and loci information were listed in **Table EV5**. Data are presented as the mean ± SD from 3 biological replicates. \**P<*0.05; n.s.: not significant.

The changes of H3K27-ac in GDC-resistant cells prompted us to determine the alterations in the chromatin occupation of BRD4, which recognizes acetylated histone such as H3K27-ac via bromodomains. BRD4 ChIP-seq analysis demonstrated that a large number of genomic loci in GDC-resistance cells lost BRD4 binding, which was consistent with the overall loss of H3K27-ac, (**Fig 3B**). However, similar to the gain of H3K27-ac, 841 genome loci gained significant BRD4 binding (**Fig 3B**). Through integrated analysis, we identified 210 loci with co-gained H3K27-ac level and BRD4 binding in GDC-resistant PC-3 cells, and 10% and 90% of these loci are located in promoter and non-promoter regions (e.g. enhancers), respectively (**Figs 3C and D, and EV3D**). The gain of H3K27-ac and BRD4 binding at representative genomic loci was confirmed by ChIP-qPCR (**Figs 3E and EV3A**). These results indicate that despite global decrease of histone acetylation and loss of BRD4 binding at a large number of gene loci, there are a subset of islanded regions co-gaining H3K27-ac level and BRD4 occupation in GDC-resistant cells versus controls.

### Both CBP/p300 and HDAC inhibitors equivalently abolish locus-specific gain of BRD4 binding in GDC-resistant cells

PI3K/AKT inhibitor resistant cells depend on the activity of BRD4 and CBP/p300 for growth (**Fig 1A-C**). These findings led us to hypothesize that—under the pressure of global decline of histone acetylation—CBP/p300 activity catalyzes acetyl modifications in critical genome loci and thereby facilitates BRD4 binding, downstream gene up-regulation and, consequently, drug resistance. To test this hypothesis, we examined whether H3K27-ac- and BRD4-gained loci in GDC-resistant cells are vulnerable to CBP/p300 inhibition. Analysis of ChIP-seq data showed that iCBP112 treatment reversed GDC-induced increases in H3K27-ac levels and BRD4 binding at the 210 gained loci in GDC-resistant PC-3 cells (**Fig 4A and B**). This result is consistent with our observation that CBP/p300 inhibitors abolish acquired PI3K/AKT inhibitor resistance (**Fig 1A-C**), emphasizing the potential connection of these loci to drug resistance. Examples of the effect of iCBP112 on specific loci are shown in **Fig 4B** and such effect was further confirmed by ChIP-qPCR at multiple loci (**Figs 4C and EV3E**).

**Figure 4.**
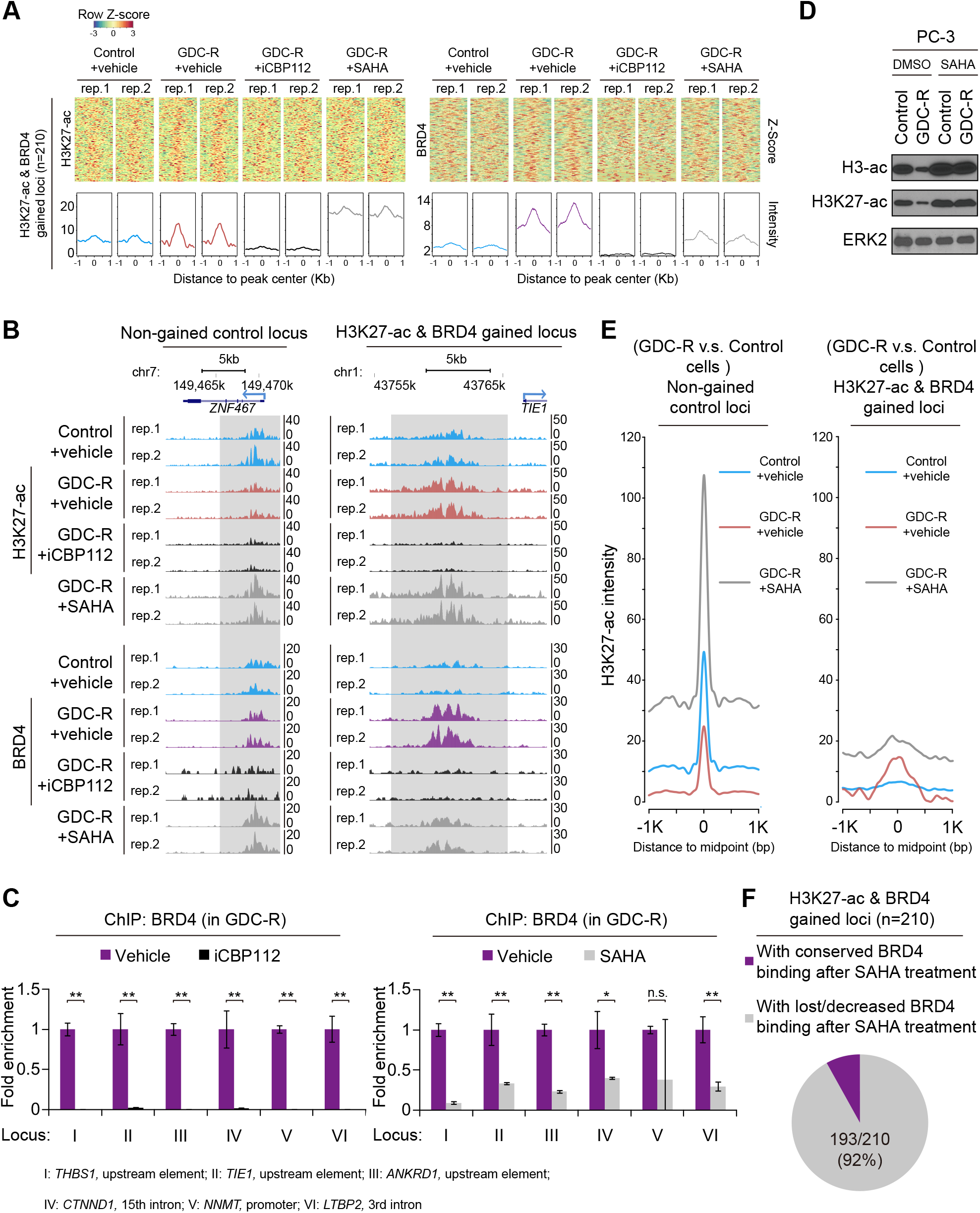
Both CBP/p300 and HDAC inhibitors equivalently abolish locus-specific gain of BRD4 binding in GDC-resistant cells. A. Z-score heatmaps show H3K27-ac and BRD4 co-gained loci in the GDC-resistant (GDC-R) cells mock treated or treated with iCBP112 or SAHA for 24 hrs. H3K27-ac ChIP-seq data was normalized with spike-in *Drosophila* chromatin DNA. Each locus is shown in a horizontal window of ±1kb from the peak center. The average intensity of all the peaks in each group was calculated and shown underneath the heatmap. B. Examples of the genome loci with or without gain of H3K27-ac and BRD4 in the PC-3 GDC-R cells. H3K27-ac ChIP-seq data was normalized with spike-in *Drosophila* chromatin DNA. Changes in these loci after iCBP112 or SAHA treatment are also shown. Gray boxes, the regions with or without gain of H3K27-ac level and BRD4 binding. rep, ChIP-seq replicates. C. ChIP-qPCR analysis of BRD4 binding at the indicated genome loci in the PC-3 GDC-R cells treated with or without iCBP112 and SAHA. Data are presented as the mean ± SD from 3 biological replicates. \**P<*0.05, \*\**P<*0.01; n.s.: not significant; duplicate ChIP experiments. D. PC-3 control and GDC-R cells were treated with DMSO or 1 μM SAHA for 24 hrs followed by western blotting. E. Meta-loci analysis of H3K27-ac intensity at H3K27-ac and BRD4 co-gained or non-gained loci in the control and GDC-R PC-3 cells treated with or without SAHA. F. Pie graph shows the percentage of the (H3K27-ac and BRD4 co-gained) loci in the GDC-R cells either with conserved BRD4 binding or with lost/decreased BRD4 binding after SAHA treatment.

GDC-induced inhibition of histone acetylation, including H3K27-ac, was completely reversed by treatment of GDC-R cells with HDAC inhibitor SAHA (vorinostat) (**Figs 4D and EV3F**). Given that HDAC inhibitors also attenuate PI3K/AKT inhibitor resistance (**Figs 1A-C**), we sought to determine how HDAC inhibitor would impact those resistance-related loci. By analyzing H3K27-ac spike in ChIP-seq and BRD4 ChIP-seq in mock versus SAHA-treated GDC-resistant PC-3 cells, we found that SAHA treatment resulted in a global increase in H3K27-ac levels in GDC-R cells; however, the increase was much lower at the H3K27-ac-gained loci (< 2 fold) compared to the regions outside these loci (> 4 fold) (**Figs 4A and E**). This result is consistent with our finding that BRD4 binding was lost or decreased at the majority (193 of 210, 92%) of the gained loci in GDC-R cells after SAHA treatment (**Figs 4A-C, 4F and EV3F**), suggesting that more BRD4 proteins are directed to the non-gained regions with massively increased H3K27-ac, which is supported by our BRD4 ChIP-seq observations (**Fig 4A and B**).

### Growth factor and receptor genes gain histone acetylation at their genomic loci and are up-regulated in drug resistant cells

Selective gain of histone acetylation at specific genomic loci in GDC-resistance cells implies their connections with up-regulation of a unique group of genes and their role in drug resistance. Through integrated analysis of RNA-seq and histone acetylation ChIP-seq data we identified a group of 123 up-regulated genes that were associated with increased levels of histone acetylation marks including H3K27-ac, H3K9/K14-ac and H4-ac at their genomic loci in GDC-resistant PC-3 cells (**Fig 5A**). We performed pathway enrichment analysis. Apart from down-regulated genes or up-regulated genes without obvious gain of histone acetylation at their genomic loci in the GDC-R cells (**Fig EV4A**), 123 genes with gained H3/H4-ac were largely enriched with growth factor/receptor (GF/R) genes that are closely related to EGFR or IGF1R signaling pathways, including *PDGFB, IGF1R* and *HBEGF* (**Fig 5B and C**).

**Figure 5.**
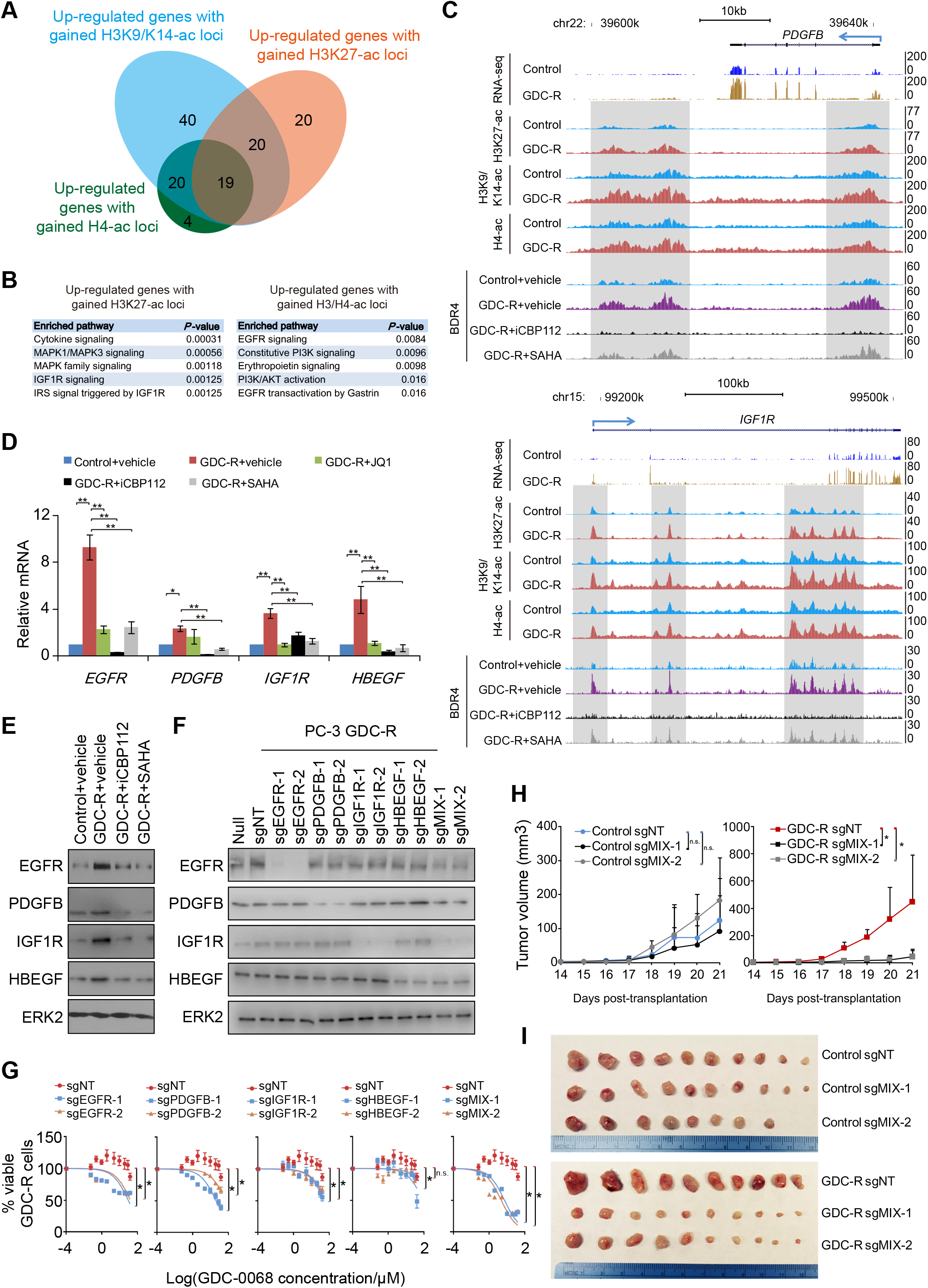
Growth factor and receptor genes gain histone acetylation at their genomic loci and are up-regulated in drug resistant cells. A. 123 transcriptionally up-regulated genes with gained histone acetylation at their genomic loci in the GDC-R PC-3 cells. B. Pathway enrichment analysis of the up-regulated genes with gained H3K27-ac or H3/H4-ac (H3K9/K14/K27-ac or H4-ac) at their genomic loci in the GDC-R PC-3 cells. C. RNA-seq and ChIP-seq profile at the indicated genomic loci in the control and GDC-R PC-3 cells treated with or without iCBP112 and SAHA. Gray boxes, the regions with gain of H3K27-ac level and BRD4 binding in the GDC-R cells. Data from two replicates were integrated. D. RT-qPCR analysis of mRNA levels of indicated genes in the control and GDC-R PC-3 cells 24 hrs after the indicated treatments. Data are presented as the mean ± SD from 3 biological replicates. \**P<*0.05, \*\**P<*0.01. E. The PC-3 control and GDC-R cells were treated as indicated for 24 hrs before analyzed by western blotting. IGF1R was detected with an antibody against the IGF1Rβ peptides. F. The PC-3 GDC-R cells were infected with lentivirus expressing non-targeting (NT) or gene-specific sgRNAs. sgMIX-1 represents pooled sgRNAs of sgEGFR-1, sgPDGFB-1, sgIGF1R-1 and sgHBEGF-1, and the same for sgMIX-2. Stably infected cells were harvested for western blotting. G. Stable cell lines generated from the GDC-R PC-3 cells as in (F) were treated with GDC-0068 for 48 hrs prior to MTS assays. The sgNT group was used as a common control for other groups. Data are presented as the mean ± SD from 3 biological replicates. \**P<*0.05; n.s.: not significant; ANOVA. H, I. Stable cell lines generated from the GDC-R PC-3 cells as in (F) were inoculated into SCID mice. Growth of tumors in each group was monitored every other day for 21 days (H) and tumors were isolated and photographed at the end of measurement (I). Data are presented as the mean ± SD from different tumors (n = 10). \**P<*0.05; n.s.: not significant; ANOVA. Loss of two tumors in the control sgMIX-2 group was due to the animal mortalities before the end point.

To further define the mechanism(s) that cause upregulation of target genes in GDC-R cells, we performed transcription factor binding motif analysis. We identified three consensus DNA motifs that are enriched at 1,332 H3K27-ac-gained peaks, but none of them matches with any transcription factor in three different databases covering approximately 850 transcription factors (**Fig EV4B and C**). Additionally, BRD4 occupancy and H3K27ac are the hallmark of super-enhancers; they are considered dominant chromatin structures that define the transcriptome program of cancerous and normal cells (Hnisz *et al*, 2013; Loven *et al*, 2013; Pelish *et al*, 2015). Since a large number of gained H3K27-ac peaks were located in enhancers (**Fig EV3C and D, and Table EV3**), we employed the ROSE program to define super-enhancers. We found that only a small portion (4 of 59, 6.7%) of up-regulated genes with gained H3K27-ac are at super-enhancer loci and are not enriched in the GF/R signaling pathways (**Figs 5B and EV4D**). These data suggest that super-enhancers may not play a major role in acquisition of GDC resistance in these cells, although our data cannot entirely rule out this possibility.

iCBP112, SAHA or JQ1 treatment decreased glucose consumption and cell proliferation, consistent with their antagonistic effects on the growth of PI3K/AKT inhibitor resistant cells (**Fig 1A-C**). These effects were mediated by G1 cell cycle arrest but not apoptosis (**Fig EV4E-I**). Reverse transcription and quantitative PCR (RT-PCR) analysis showed that iCBP112 or SAHA treatment also blocked the upregulation of GF/R genes in GDC-R cells (**Fig 5D and E**). The correlation of iCBP112 or SAHA-induced suppression of these genes and the ablation of AKT inhibitor resistance suggests that these GF/R genes might contribute to AKT inhibitor resistance.

To directly test whether upregulation of GF/R genes plays a causal role in GDC resistance, control and GDC-resistant PC-3 cells were transfected with lentivirus expressing CRISPR/Cas9 and gene-specific small guide RNAs (sgRNAs) for *EGFR, PDGFB, IGF1R* and *HBGF*. Depletion of these GF/R genes individually slightly, but depletion together largely re-sensitized GDC-resistant PC-3 cells to GDC in vitro and in mice (**Fig 5F-I**). Similar to the effect of iCBP112 and SAHA, depletion of these GF/R genes induced G1 cell cycle arrest (**Fig EV4J and K**). These results demonstrated that upregulation of a subset of GF/R genes is important for proliferation of GDC-R PC-3 cells.

A recent study reports that rapid loss of histone acetylation induces differentiation of embryonic stem cells (Moussaieff *et al.*, 2015). We speculated that loss of histone acetylation caused by metabolic stress might trigger expression of resistance-related GF/R genes. Consistent with the finding that blockage of glycolysis by 2-DG reduced global histone acetylation in both PC-3 and C4-2 cells (**Fig 2G**), 2-DG treatment also increased GF/R gene expression; however, this effect was blocked by pre-treatment with SAHA (**Fig EV5A and B**). These results indicated that global loss of histone acetylation is one of the factors that are essential for the upregulation of GF/R genes in GDC-resistant cells. In PTEN-positive cells such as LAPC-4 and VCaP, GDC treatment only robustly decreased histone acetylation in the presence of EGF (**Fig EV5C**). Unlike PTEN-negative (i.e. constitutively active AKT) cells (**Fig 1B**), EGF-stimulated PTEN-proficient (i.e. transiently active AKT) cells were more sensitive to GDC but more resistant to iCBP112 or SAHA-induced suppression of cell growth and expression of GDC-resistant genes such as *IGF1R* and *PDGFB* compared to mock-treated cells (**Fig EV5D and E**). Thus, it is conceivable that oncogene addiction due to the constitutive activation of the PI3K pathway in PTEN-null cells could epigenetically prime the resistant gene loci to respond to both bromodomain and HDAC inhibitors.

### BET and CBP/p300 bromodomain dual inhibitor NEO2734 and butyrate are both active in PI3K/AKT inhibitor-resistant cancer cells

Bromodomain proteins BRD4 and CBP/p300 co-operate to activate transcription. Given that individual inhibition of BRD4 and CBP/p300 antagonizes PI3K/AKT inhibitor resistance (**Fig 1A-C**), we sought to determine whether the combination of these two types of inhibitors results in synergy. Co-treatment of JQ1 and CBP/p300 inhibitor CPI-637 largely inhibited the growth of GDC-resistant PC-3 cells in vitro and in vivo (**Fig EV6A and B**). Consistent with its effects on GDC-resistant cells in culture (**Fig EV6A**), SAHA also significantly inhibited growth of the resistant cells in vivo (**Fig EV6B**). The anti-tumor effect of CBP/p300 inhibitors did not appear to be caused by DNA damage (**Fig EV6C and D**). While SAHA or bromodomain inhibitor alone can effectively inhibit GDC-resistant cell growth at high doses, the combination of SAHA with JQ1 or iCBP112 resulted in synergistic inhibition of GDC-resistant cell growth at lower doses (**Fig EV6E and F**), which is consistent with the findings in other cancer types such as melanoma (Heinemann *et al*, 2015).

NEO2734 has been recently demonstrated as a multi-bromodomain inhibitor which targets BRD2, BRD3, BRD4, CBP and p300 (Yan *et al*, 2019). We examined the efficacy of NEO2734 in overcoming PI3K/AKT resistance. Butyrate, a four-carbon short-chain fatty acid, is a natural component of the colonic milieu and acts as a HDAC inhibitor to inhibit cancer cell growth at concentrations detectable in the mammalian colon (Archer *et al*, 1998; McIntyre *et al*, 1993; McIntyre *et al*, 1991). We also examined the inhibitory effect of this naturally occurring HDAC inhibitor on PI3K/AKT inhibitor resistance. We demonstrated that GDC-resistant PC-3 cells were more sensitive to both NEO2734 and butyrate than the control cells in vitro (**Fig 6A and B**). Although butyrate is effective only at mM concentrations in vitro (**Fig 6B**), it is more specific against growth of the GDC-resistant tumors than that of control tumors in mice (**Fig 6C and D**). RT-qPCR analysis confirmed that mRNA expression of GF/R genes such as *PDGFB* and *IGF1R* was increased in the GDC-resistant xenografts compared to the control tumors, but was reversed by either NEO2734 or butyrate (**Fig 6E**). These data imply an important role of GF/R genes in development of AKT inhibitor resistance in vivo. We also assessed the vulnerability of different PI3K/AKT inhibitor resistant cell lines to NEO2734 and butyrate. Most, especially GDC-resistant cell lines, were hypersensitive to both inhibitors (**Fig 6F and G**). Together, these results posit new therapeutic options to manage PI3K/AKT inhibitor resistance at the chromatin level rather than targeting individual genes. These findings further confirm the efficacy of both bromodomain and HDAC inhibitors in treating PI3K/AKT inhibitor resistant cells.

**Figure 6.**
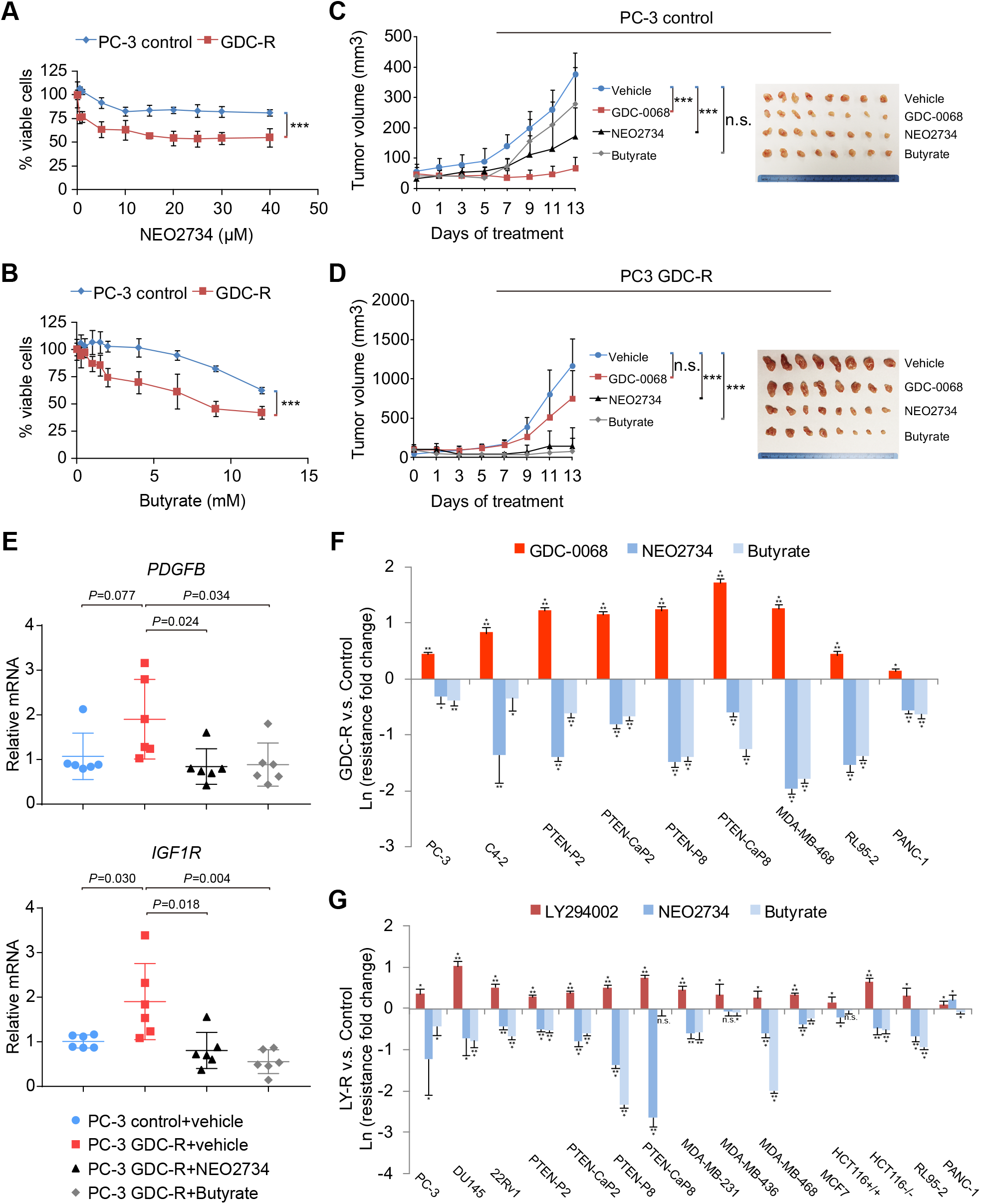
BET and CBP/p300 bromodomain dual inhibitor NEO2734 and butyrate are both active in PI3K/AKT inhibitor-resistant cancer cells. A, B. PC-3 control and GDC-R cells were treated with NEO2734 (A) or butyrate (B) for 48 hrs before MTS assays. Data are presented as the mean ± SD from 3 biological replicates. \*\*\**P<*0.001; ANOVA. C, D. PC-3 control and GDC-R cells were inoculated in SCID mice and tumor growth was monitored and treated every other day with vehicle, GDC-0068 (100 mg/kg), NEO2734 (10 mg/kg) or butyrate (1,000 mg/kg) (Left). Tumors were isolated and photographed at the end of measurement (Right). Data are presented as the mean ± SD from replicates (n = 8). \*\*\**P<*0.001; n.s.: not significant; ANOVA. E. RT-qPCR analysis of the GF/R genes *PDGFB* and *IGF1R* in six randomly collected xenografts (n = 6) from different groups shown in (C) and (D). Data are presented as the mean ± SD from different tumors (n = 6). The exact *P* values are indicated. F, G. GDC-0068 (or LY294002) -and mock-trained cells were treated with GDC-0068 (or LY294002), NEO2734 or butyrate (at the IC50 concentration of the compound in the control cells) for 48 hrs. The resistance of each compound was calculated as described in **Fig 1B and C**. Data are presented as the mean ± SD from 3 biological replicates. \**P<*0.05, \*\**P<*0.01, \*\*\**P<*0.001; n.s.: not significant.

### Enhanced binding of CBP/p300 and inadequate occupancy of HDACs on chromatin permit locus-specific gain of H3K27-ac upon AKT inhibition

Next, we sought to define what could be the major mechanism underlying the nearly equivalent effects of bromodomain and HDAC inhibitors on BRD4-regulated gene expression and PI3K/AKT inhibitor resistance.

Observing that the gain of H3K27-ac level and BRD4 binding at resistance-related loci can be reversed by CBP/p300 inhibitors, we reasoned that these loci are likely occupied by acetyltransferases CBP and p300 in the resistant cells. They would thus retain persistent histone acetylation under acetyl-CoA stress. To test this hypothesis, we performed CBP and p300 ChIP-seq and found that both CBP and p300 enrichment and the levels of other histone acetylation marks including H3K9/K14-ac and H4-ac were higher at the H3K27-ac gained loci in GDC-R cells compared to control cells (**Fig 7A and B**).

**Figure 7.**
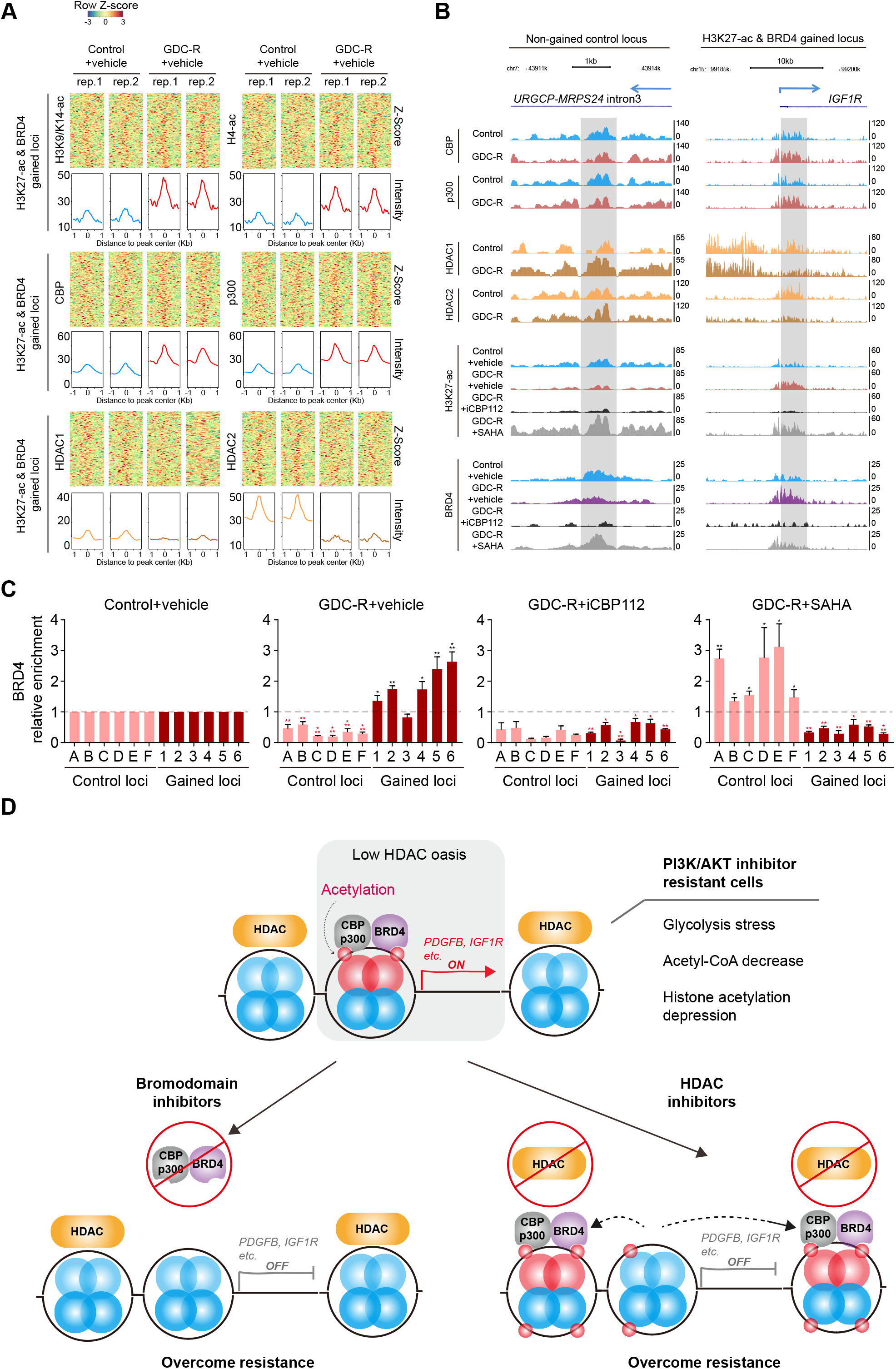
Enhanced binding of CBP/p300 and inadequate occupancy of HDACs on chromatin permit locus-specific gain of H3K27-ac upon AKT inhibition. A. Z-score heatmaps of H3K9/K14-ac, H4-ac, CBP, p300, HDAC1 and HDAC2 ChIP-seq signals at the (both H3K27-ac and BRD4) gained loci in the GDC-resistant PC-3 cells. Each locus was shown in a horizontal window of ±1kb from the peak center. The average intensity of all the peaks in each group was calculated and shown underneath the heatmap. B. ChIP-seq profile for the indicated proteins and H3K27-ac at the (H3K27-ac and BRD4) co-gained and non-gained loci in the control and GDC-R PC-3 cells treated with or without iCBP112 and SAHA. Gray boxes, the regions of interest with or without gain of H3K27-ac level and BRD4 binding in the GDC-R cells. Data from two replicates were integrated. C. ChIP-qPCR analysis of BRD4 binding at the gained and non-gained control loci in the control and GDC-R PC-3 cells with or without iCBP112 and SAHA treatment. Primers and loci information were listed in **Table EV5**. Gained loci include 1: *EGFR*, upstream element; 2: *HBEGF*, downstream element; 3: *TGFB2*, upstream element; 4: *IGF1R*, near-promoter element; 5: *PDGFB*, downstream element; 6: *ZNF365*, upstream element. Dashed line indicated the normalized BRD4 fold enrichment in the control group. Data are presented as the mean ± SD from 3 biological replicates. \*\**P<*0.01, \*\*\**P<*0.001. D. A hypothetical working model of the current study. Top: PI3K/AKT inhibitors induce glycolysis crisis, shortage of acetyl-CoA supply and global loss of histone acetylation including H3K27-ac, a stressful condition but being accompanied by increased H3K27-ac and BRD4 binding at some CBP/p300 enriched, HDAC-deficient genomic loci (low HDAC ‘oases’) such as GF/R gene regions and ultimately resistance to PI3K/AKT inhibitors. Bottom Left: Bromodomain inhibitors directly block the binding of CBP/p300 and BRD4 at the HDAC low-occupied GF/R gene loci, abolishing expression of GF/R genes and PI3K/AKT inhibitor resistance. Bottom Right: While treatment of HDAC inhibitors induces a massive global increase in histone acetylation, the increase is much lower at the HDAC low-occupied GF/R gene loci than that in the rest of the genome, thereby resulting in more CBP/p300 and BRD4 protein binding to the regions beyond the GF/R loci, decreased expression of GF/R genes, and overcoming of PI3K/AKT inhibitor resistance.

In GDC-resistant cells, we observed that HDAC inhibitor had a much smaller effect on elevation of H3K27-ac at gained loci compared non-gained loci (**Fig 4E**). This observation prompted us to hypothesize that the drastic gain of H3K27-ac and BRD4 binding might be due to low or no HDAC binding at the gained loci in the resistant cells. We carried out ChIP-seq for HDAC1 and HDAC2 (HDAC3 was excluded due to lack of suitable antibody for ChIP) in both control and GDC-R PC-3 cells. In support with our hypothesis, we confirmed that occupation of HDAC, especially HDAC2 proteins at the H3K27-ac and BRD4 co-gained loci was lower in resistant cells compared to control cells (**Fig 7A and B)**.

We found that the basal H3K27-ac level in non-gained control loci was higher than that in the gained loci; however, after long-term treatment of GDC the H3K27-ac level at control loci was downregulated whereas the H3K27-ac level at gained loci was upregulated (**Fig 4E**). Importantly, CBP/p300 binding was higher, but HDAC occupancy was lower at the gained loci in the GDC-R cells compared to control cells (**Fig 7A and B**). These results provide a plausible explanation to our observation that iCBP112 treatment decreased H3K27-ac levels at gained loci in GDC-resistant cells (**Fig 4A**) and that SAHA treatment resulted in a marked increase in the H3K27-ac levels at control loci with minimal effect at gained loci (**Fig 4E**).

Next, we examined BRD4 binding at CBP/p300 enriched, HDAC-depleted loci under different cellular conditions. We found that the BRD4 binding was diminished at non-gained loci, but enhanced at gained loci in GDC-resistant cells (**Fig 7C**). iCBP112 treatment blocked the increased binding of BRD4 at gained loci (**Fig 7C**). SAHA treatment increased BRD4 binding in non-gained loci but decreased BRD4 binding at gained loci (**Fig 7C**, second versus fourth panels). The differences in BRD4 binding between control and GDC-R cells were not caused by BRD4 expression level changes (**Fig EV7**)

## Discussion

Bromodomain and HDAC inhibitors are two classes of compounds with distinct molecular mechanism of action although both are closely related to histone acetylation (Manzotti *et al*, 2019). In various cancer cell line models we demonstrate that PI3K/AKT inhibitors induce suppression of glycolysis, down-regulation of acetyl-CoA supply and, ultimately, a significant decrease in histone acetylation including H3K27-ac. We found that histone acetylation (especially H3K27-ac) level is unexpectedly elevated in a subset of GF/R gene loci despite glycolytic stress, acetyl-CoA shortage and global depression of histone in company with increased GF/R gene expression. Others have shown that expression of the GF/R gene *IGF1R* is important for the survival of PI3K inhibitor resistant cancer cells (Chandarlapaty *et al.*, 2011; Stratikopoulos *et al.*, 2015). We extend these observations by showing that in addition to IGF1R, cooperation with other GF/R factors such as EGFR, PDGFB and HBEGF is also important for development of AKT inhibitor resistance. Mechanistically, we discovered that almost all these GF/R gene loci have increased CBP/p300 binding and decreased occupancy of HDAC proteins in resistant cells. Our findings support the notion that abundant CBP/p300 binding together with sparse HDAC occupation at GF/R gene regions epigenetically prime these genomic loci for a surge of histone acetylation such as H3K27-ac when cells confront glycolysis stress, acetyl-CoA shortage and globally decreased histone acetylation caused by PI3K/AKT inhibitors (**Fig 7D**, upper). Intriguingly, increased H3K27-ac level and BRD4 occupancy at these loci are equivalently suppressed by both bromodomain and HDAC inhibitors. The finding that two types of molecules with distinct modes of action exert the similar inhibitory effect on the expression of almost all the GF/R genes we identified provides a plausible explanation to the effective killing of the PI3K/AKT inhibitor-resistant cancer cells by both bromodomain and HDAC inhibitors.

Inadequate HDAC occupancy at GF/R gene loci appears to be a critical molecular trait of the PI3K/AKT inhibitor resistant cancer cells. We observed that although resistance-associated gene loci are mostly HDAC binding-deficient, not all HDAC-inadequate loci could be activated after AKT inhibition. Our data suggest that deficient HDAC binding is necessary but not sufficient to activate these GF/R genes. It is conceivable that sequence specific transcription factors might play a key role in transcriptional activation of the GF/R genes identified. Surprisingly, no binding motif of specific transcription factors was identified through DNA binding motif analysis. Our results suggest that the increased expression of GF/R genes in PI3K/AKT inhibitor resistant cells is likely driven by a diverse pool of transcription factors.

Our findings also support the notion of complex correlation between transcription factor binding and target gene expression. It has been shown previously that different types of cancer cells may acquire PI3K/AKT inhibitor resistance through induction of different sets of genes mediated by different transcription factors such as FOXO, ER and AR (Carver *et al.*, 2011; Chandarlapaty *et al.*, 2011; Garrett *et al.*, 2011; Mulholland *et al.*, 2011; Toska *et al.*, 2017). It seems very difficult to predict the resistant genes through their DNA sequence. However, regardless of which GF/R genes can be activated in resistant cancer cells, our study demonstrates that reversing resistance at chromatin level offers a general solution. Given that treatment of PI3K/AKT inhibitors inevitably leads to suppression of glycolysis and acetyl-CoA shortage, it is not surprising that only those HDAC binding-deficient gene loci with locally enhanced CBP/p300 activity can maintain or even boost local histone acetylation levels. Thus, the locally enhanced CBP/p300-BRD4 activity at GF/R gene loci represents an Achilles heel for PI3K/AKT inhibitor-resistant cells, making it susceptible to both BRD4 and CBP/p300 bromodomain inhibitors individually (**Fig 7D**, lower left). On the other hand, we demonstrate that deficient HDAC binding at GF/R gene loci is an additional vulnerability in these resistant cells. Given low HDAC occupancy at these loci, it is not surprising that HDAC inhibitors struggle to increase histone acetylation at these genomic regions but rather cause massive increases in histone acetylation outside these regions. As a result, HDAC inhibitor treatment may generate a panoply of bromodomain protein binding sites to attract BRD4 (and potentially CBP/p300) (Dey *et al*, 2003) away from the GF/R gene loci, ultimately leading to down-regulation of GF/R gene expression and overcoming resistance to PI3K/AKT inhibitors (**Fig 7D**, lower right). Thus, our findings uncover a new mechanism underlying HDAC inhibitor-induced redistribution of BRD4 among drug resistance gene loci in stressed cells, thereby extending the finding of a previous report that HDAC inhibitor treatment induces redistribution of BRD4 from promoters to gene bodies and thereby facilitates gene transcription elongation in unstressed cells (Greer *et al*, 2015).

Our study provides a new molecular insight into the mechanism of action of bromodomain and HDAC inhibitors. Bromodomain inhibitors are generally competitive inhibitors that block the interaction between bromodomain (present in ‘reader’ or ‘reader/writer’ proteins) and acetyl-lysine residues on histones. Since the discovery of the first generation BET bromodomain inhibitors such as JQ1 and I-BET in the recent decade (Filippakopoulos *et al*, 2010; Nicodeme *et al*, 2010), it has been shown that bromodomain inhibitors have potent anti-cancer activity in leukemia and solid tumors (Zhang *et al*, 2017). In contrast, HDAC inhibitors target the ‘erasers’ of histone acetylation. Both natural and synthetic HDAC inhibitors have been validated as anti-cancer agents and some of them have been approved for clinical use (Grant *et al*, 2007; Yoshida *et al*, 1990). Interestingly, bromodomain and HDAC inhibitors both inhibit the growth of cancer cells significantly despite their entirely opposite modes of action (Heinemann *et al.*, 2015). Our data demonstrate that under certain conditions—such as global loss of histone acetylation caused by PI3K/AKT inhibitors—these two distinct classes of inhibitors may target the same set of gene loci. Based on the model we proposed (**Fig 7D**), both bromodomain and HDAC inhibitors are potent suppressors of growth-promoting gene expression in genomic regions with high occupancy of bromodomain proteins and low occupancy of HDACs.

In summary, our study uncovers a histone acetylation-centric program that conveys PI3K/AKT inhibitor resistance. We demonstrate that PI3K/AKT inhibitors invariably induce glycolysis stress, shortage of acetyl-CoA supply and ultimately inhibition of histone acetylation. We further show that, despite these stressful conditions, H3K27-ac level and BRD4 binding are up-regulated in the loci of a subset of GF/R genes that are essential for the survival of PI3K/AKT resistant cells. We provide evidence that both bromodomain and HDAC inhibitors can collapse this epigenetic program. Our findings imply that pharmaceutical intervention at the chromatin level could be a promising strategy to circumvent PI3K/AKT inhibitor resistance, thereby offering two ‘silver bullets’ to overcome this obstacle in cancer therapy.

## Materials and Methods

### Antibodies and reagents

Antibodies used are as follows: AKT (Cat# 9272), p-AKT(S473) (Cat# 9271), mTOR (Cat# 2983), p-mTOR(S2448) (Cat# 5536), GSK-3β (Cat# 9315), p-GSK-3β(S9) (Cat# 9322), H3 (Cat# 9715), H3K27-me3 (Cat# 9733), H3K14-ac (Cat# 7627), H3K56-ac (Cat# 4243), H4K5-ac (Cat# 8647), H4K12-ac (Cat# 13944), H4K16-ac (Cat# 13534), CBP (Cat# 7425), p300 (Cat# 54062), HK2 (Cat# 2867), IGF1Rβ (Cat# 3027) purchased from Cell Signaling Technology; H3-ac (Cat# ab47915), H3K9/K14-ac (Cat# ab232952), H3K27-ac (Cat# ab4729), H3K9-ac (Cat# ab4441), H3K18-ac (Cat# ab1191), H3K23-ac (Cat# ab177275), H4K8-ac (Cat# ab45166), BRD4 (Cat# ab128874), HDAC1 (Cat# ab7028), HDAC2 (Cat# ab7029), FBP1 (Cat#ab109732) purchased from Abcam; H4 (Cat# sc-25260), GAPDH (Cat# sc-137179), ERK2 (Cat# sc-1647), EGFR (Cat# sc-03), PDGFB (Cat# sc-365805), HBEGF (Cat# sc-365182) purchased from Santa Cruz Biotechnology; H4-ac (K5/K8/K12-ac) (Cat# PA5-40083) purchased from Thermo Fisher.

Chemical and other reagents used are as follows: Paclitaxel (Taxol) (Cat# T7402), Trichostatin A (TSA) (Cat# T8552), Sodium butyrate (Butyrate) (Cat# B5887), Flavopiridol HCl (Cat# F3055), Recombinant human EGF (Cat# E9644) and Glucose Quantification Kit (Cat# GAGO-20) purchased from Sigma-Aldrich; Cisplatin (Cat# 232120) purchased from Calbiochem; Camptothecine (CPT) (Cat# 159732) purchased from MP Biomedicals; GSK1120212 (Cat# A-1258) purchased from Active Biochemicals; SB202190 (Cat# 21201), iCBP112 (Cat# 14468), Vorinostat (SAHA) (Cat# 10009929) and 2-Deoxy-D-glucose (2-DG) (Cat# 14325) purchased from Cayman; MLN4924 (Cat# 505477), JSH23 (Cat# 481408) and Recombinant human IGF-1 (Cat# GF138) purchased from Millipore; JQ1 (Cat# M2167) and RAD001 (Cat# M1709) purchased from AbMoleBioScience; GSK126 (Cat# T2079) purchased from TargetMol; Recombinant human bFGF (Cat# 13256029) purchased from Gibco; LY294002 (Cat# PHZ1144) purchased from Invitrogen; MK-2206 (Cat# S1078), Ku-55933 (Cat# S1092), PD-0332991 (Cat# S1116), Etoposide (ETO) (Cat# S1225), Olaparib (Cat# S1060), QNZ (Cat# S4902) and EPZ-6438 (Cat# S7128) purchased from Selleckchem; GDC-0068 (Cat# HY-15186A), CPI-637 (Cat# HY-100482) and RGFP966 (Cat# HY-13909) purchased from MedChem Express; Acetyl-CoA Quantification Kit (Cat# K317-100) purchased from BioVision.

### Cell lines, cell culture and transfection

Human cancer cell lines were obtained from the American Type Culture Collection (ATCC). PC-3, C4-2, 22Rv1 and PANC-1 cell lines were cultured in the RPMI 1640 medium supplemented with 10% fetal bovine serum (FBS). DU145, VCaP, LAPC-4, MCF7 and RL95-2 cell lines were maintained in the Dulbecco’s modified Eagle’s medium (DMEM) supplemented with 10% FBS. MDA-MB-231, MDA-MB-436 and MDA-MB-468 cells were maintained in the Leibovitz’s L-15 medium supplemented with 10% FBS. HCT116 Tp53 +/+ (HCT116 +/+) and HCT116 -/- cells were maintained in the McCoy’s 5A medium supplemented with 10% FBS. Murine prostate cancer cell lines PTEN-P2, PTEN-CaP2, PTEN-P8 and PTEN-CaP8 were kindly provided by Dr. Hong Wu (Jiao *et al*, 2007) and were maintained in DMEM supplemented with 10% FBS, 25 μg/mL bovine pituitary extract, 5 μg/mL bovine insulin and 6 ng/mL recombinant human epidermal growth factor. These cells were genotyped by standard genomic PCR techniques as indicated in the previous report (Jiao *et al.*, 2007). HEK293T cells were purchased from ATCC and cultured in DMEM supplemented with 10% FBS. See also **Table EV2** and references (Deer *et al*, 2010; Tate *et al*, 2019) for additional cell line information. All cell lines were cultured at 37°C with 5% CO_2_. *Drosophila* S2 cells (purchased from ATCC, catalog number CRL-1963) were cultured in Schneider’s *Drosophila* media (Life Technologies, catalog number 21720-024) supplemented with 10% FBS and cultured at 21°C without additional CO_2_. Plasmocin (Invivogen) was added to cell culture media to prevent mycoplasma contamination. Mycoplasma contamination was tested monthly using the Lookout Mycoplasma PCR Detection Kit (Sigma-Aldrich). All cells were mycoplasma-free. Transfection was performed using Polyethylenimine (PEI, 25kD linear, Polysciences).

### Comparison of the effects of HDAC inhibitors and BRD4 knockdown on the transcriptome in different cancer cell lines

The Cell Connectivity Map (CMAP) is a compendium of gene expression signatures induced by chemical compounds or genetic perturbations (termed perturbagens) in multiple cell lines. For each small molecule treatment or single gene knockdown/over-expression in a specific cell line, the L1000 assay platform measured the reduced representation of total transcriptome (∼1,000 land mark transcripts), recorded as an expression signature (Lamb *et al.*, 2006; Subramanian *et al.*, 2017). CMAP has collected approximately 1.3 million expression signatures of over 8,000 perturbagens in at least 7 cell types, and the “distance” between any two expression signatures is quantified using Kolmogorov-Smirnov test (K-S test). More similar expression signatures showed “shorter distance” and related higher “connectivity value”., We searched CMAP (https://clue.io/) using the command: /conn “mocetinostat” “NCH-51” “panobinostat” “scriptaid” “THM-I-94” “vorinostat” “trichostatin-a” or the command: /conn “BRD4-KD”. The resultant connectivity data was then downloaded and visualized as heatmaps.

### Multi-bromodomain inhibitor NEO2734

NEO2734 (also known as EP31670) was identified from a drug discovery collaboration between the University of Miami, Epigenetix Inc and the Neomed Institute (Yan *et al.*, 2019). The activity of NEO2734 in inhibition of the bromodomains in BET and CBP/p300 was initially tested with the BROMOscan platform (DiscoverX/Eurofins). The EC50 of NEO2734 has been demonstrated at nanomolar levels in specific inhibition of all bromodomains in BRD2, BRD3, BRD4, CBP and p300 proteins.

### Lentivirus infection

shRNAs targeting human *FBP1* mRNA were constructed into the pLKO-based gene-knockdown lentiviral vector. sgRNAs targeting human *EGFR, PDGFB, IGF1R* or *HBEGF* genes were inserted into the pLenti-crispr-V2 vector. Gene-specific targeting vector along with the packing constructs were transfected into HEK293T cells. Virus-containing supernatant was collected 48 hrs after transfection. Control and GDC-0068 resistant (GDC-R) PC-3 cells were infected with virus-containing supernatant in the presence of polybrene (5μg/ml) and were selected in growth medium containing 1.5μg/ml puromycin. Sequences of gene-specific shRNAs and sgRNAs are listed in **Table EV5**.

### Generation and treatment of AKT inhibitor resistant cancer xenografts in mice

Six-week-old SCID mice were generated in house and used for animal experiments. The animal study was approved by the IACUC at Mayo Clinic. All mice were housed in standard conditions with a 12 hrs light/dark cycle. 2×10^6^ PC-3 control and GDC-R cells (or lentivirus transfected stable lines based on these cells) were injected subcutaneously into the right flank of the mice. After xenografts reached an approximate size of 100 mm^3^, vehicle (10% β-cyclodextrin (intraperitoneal injection) and saline (oral administration)), GDC-0068 (100 mg per kg body weight (oral administration)), JQ1 (25 mg per kg body weight (intraperitoneal injection)) plus CPI-637 (5 mg per kg body weight (intraperitoneal injection)), SAHA (100 mg per kg body weight (oral administration)), NEO2734 (10 mg per kg body weight (intraperitoneal injection)) and sodium butyrate (1,000 mg per kg body weight (intraperitoneal injection)) were administrated individually every other day. The volume of xenografts was measured every other day for 21 days or until the largest tumor had reached approximately 1,000 mm^3^. The volume of an in vivo tumor was estimated using the formula 0.5×Length×Width^2^. Upon completion of measurement, tumor grafts were harvested and the actual volume of the ex vivo tumor was measured. The ratio of the average actual volume to the average estimated volume of the same tumors at the end point was calculated as the adjustment coefficient.

### Cell proliferation

Cell proliferation was measured via the MTS assay. Cells (2,000 per well) were seeded in 96-well plates with 100 mL of culture medium. Each well was added with 20 mL of CellTiter 96R AQueous One Solution Reagent (Promega) and absorbance was measured in a microplate reader at 490 nm. For cell growth assays, measurements were carried out at different time points. For compound sensitivity (IC50) assays, cell growth under chemical gradients was measured 48 h after treatment.

### Glucose quantification

Glucose measurements in the cell culture medium were performed using the Glucose Assay Kit (Sigma-Aldrich, GAGO-20). In brief, glucose was oxidized to gluconic acid and H_2_O_2_ by glucose oxidase. H_2_O_2_ oxidized o-dianisidine catalyzed by peroxidase. Oxidized o-dianisidine reacted with H_2_SO_4_ to form a stable colored product. The intensity of the color measured at 540 nm was proportionate to the original glucose quantity.

### Acetyl-CoA quantification

Acetyl-CoA was quantified using the PicoProbe™ Acetyl CoA Fluorometric Assay Kit (BioVision, K317-100). In brief, 2×10^6^ cells were homogenized using a Dounce homogenizer on ice. Samples were deproteinized via ice cold perchloric acid and neutralized by KOH. Free CoA in the samples was quenched before Acetyl-CoA was converted to CoA. The CoA was reacted to form NADH which interacts with PicoProbe to generate fluorescence (Ex/Em =535/587 nm). The intensity of the fluorescence was proportionate to the original acetyl-CoA quantity.

### Histone extraction

Cells were collected and washed with ice cold PBS before they were lysed with 0.5% Triton X 100 (v/v) for 10 min on ice. The cell nuclei pellet was re-suspended in 0.2 N HCl overnight at 4°C. The acid extracted sample was centrifuged and supernatant was reserved after protein quantification.

### Western blotting

Cell lysates or acid extracted histones were separated by sodium dodecyl sulfate polyacrylamide gel electrophoresis (SDS-PAGE) and transferred onto nitrocellulose membranes. The membranes were incubated with primary antibody overnight at 4°C and subsequently probed with horseradish peroxidase (HRP)-conjugated secondary antibodies. The immunoreactive blots were visualized using a chemiluminescence reagent (Thermo Fisher Scientific).

### RNA-seq and data analysis

RNA was isolated from control and GDC-RPC-3 cells using the RNeasy PlusMini Kit (Qiagen). High-quality (Agilent Bioanalyzer RIN>7.0) total RNA (riboRNA-depleted) was employed in the preparation of sequencing libraries. Samples with biological triplicates were sequenced using the Illumina HiSeq3000 platform. Genome-wide coverage signals were represented in BigWig format to facilitate convenient visualization using the UCSC Genome Browser. Gene expression was measured using RPKM (reads per kilobase exon per million mapped reads). EdgeR was used to identify differentially expressed genes. Pathway enrichment analysis was performed at the Reactome database (https://reactome.org/).

### RT-Quantitative PCR

Total RNA was extracted from tissues or cultured cells using Trizol reagent, and was reverse transcribed by using Superscript reverse transcriptase (Thermo Fisher Scientific) according to the manufacturer’s protocol. Quantitative real-time PCR (qPCR) was carried out by using SYBR Green Supermix kit with the iCycler QTX detection system (Bio-Rad). Primers are listed in **Table EV5**. The specificity of PCR product was confirmed by melting curve analysis and gel electrophoresis. All gene expression levels were normalized to that of the housekeeping gene beta-2-microglobulin (β2-MG).

### Chromatin immunoprecipitation sequencing (ChIP-seq) and ChIP-qPCR

Control and GDC-R PC-3 cells with or without iCBP112 (or SAHA) treatment were harvested for ChIP assay using validated HDAC1, HDAC2, CBP, p300, H3K9/K14-ac, H3K27-ac, H4K5/K8/K12-ac (H4-ac) and BRD4 antibodies. Experiments in two biological replicates (ENCODE standard) were performed for each group. For HDAC1, HDAC2, CBP and p300 ChIP, a double crosslink method was used to stabilize protein-chromatin interactions. Specifically, cells were first fixed with 2 mM DSG (Pierce) for 45 min prior to formaldehyde fixation. For each spike-in H3K27-ac ChIP experiment, a 20:1 ratio of human cells versus *Drosophila* S2 cells was used. S2 cells were added to human cells at the beginning of the ChIP workflow. Once *Drosophila* S2 and human cells were combined, the sample was treated as a single ChIP-seq sample throughout the experiment until completion of DNA sequencing. The input and ChIP DNA were used for preparation of sequencing libraries according to the manufacturer’s instructions for the Illumina Genome Analyzer. After PCR pre-amplification, size selection was performed and DNA fragments were in the range of 150-250 bp. The total DNA of each sample was normalized after amplification. The purified libraries were sequenced on an Illumina Hiseq 3000 Genome Analyzer. Reads were aligned using Bowtie2 against a “meta-genome” that combines the human hg19 genome and the *Drosophila* dm6 genome. Reads mapped to the human genome were normalized using a method described previously (Orlando *et al*, 2014) and reads mapped to one or two locations were included in further analysis. Peak calling was performed by MACS2 (2.1.1) with a *P*-value threshold of 1×10^−5^. BigWig files were generated for visualization via the UCSC Genome Browser. GREAT was applied to assign peaks to their potential target genes. For GDC-R cell specific peaks (and the peaks associated with genes up-regulated in GDC-R cells), the 500-bp genomic sequences surrounding each peak were extracted for MEME motif over-representation analysis with default parameters. The strongest log-odds matrix from the MEME output was shown in **Fig EV4B**. The input and ChIP DNA were then subjected to qPCR using binding-site specific primers which are listed in **Table EV5**.

### Statistical analysis

Unless otherwise noted, each in vitro experiment was repeated three times or more. In RT-qPCR, relative gene expression levels normalized by β2-MG were calculated by the formula 2^-ΔCt^, where ΔCt (critical threshold) = Ct of genes of interest-Ct of β2-MG. In ChIP-qPCR, the enrichment for specific locus was normalized by input in the same way. Fold changes of gene expression level or ChIP enrichment were calculated by the 2^-ΔΔCt^ method. The data in histograms were presented as the mean ± standard deviation. The significance of the differences was determined by Student *t* test or ANOVA. Statistical analyses were performed in GraphPad Prism software. *P*< 0.05 was considered statistically significant.

## Acknowledgements

The authors would like to thank Dr. Taro Hitosugi for his advice for measurement of acetyl-CoA. This work was supported by the Mayo Clinic Foundation (to H.H.).

## Author contributions

H.H., D.W., S.R. conceived the study. D.W., Y.Y., Y.X., D.W. generated reagents and performed experiments, D.W. performed data collection and analysis. L.W., T.W., Z.Y., J.J.O. performed data mining and NGS data analysis. Y.P. assisted D.W. to complete the mouse experiments. H.H., S.R., J.J.O. wrote the manuscript.

